# Mitochondrial respiration is required to provide amino acids during fermentative proliferation of fission yeast

**DOI:** 10.1101/2020.02.12.946111

**Authors:** Michal Malecki, Stephan Kamrad, Markus Ralser, Jürg Bähler

**Author notes:** Correspondence (MM); (JB).

## Abstract

When glucose is available, many organisms repress mitochondrial respiration in favour of aerobic glycolysis, or fermentation in yeast, that suffices for ATP production. Fission yeast cells, however, rely partially on respiration for rapid proliferation under fermentative conditions. Here we determined the limiting factors that require respiratory function during fermentation. When the electron transport chain was inhibited, supplementation with arginine was necessary and sufficient to restore rapid cell proliferation. Accordingly, a systematic screen for mutants growing poorly without arginine identified not only mutants defective in arginine synthesis but also mutants defective in mitochondrial oxidative metabolism. Genetic or pharmacological inhibition of respiration triggered a drop in intracellular levels of arginine and amino acids derived from the Krebs-cycle metabolite alpha-ketoglutarate: glutamine, lysine and glutamic acid. Conversion of arginine into these amino acids was required for rapid proliferation when the respiratory chain was blocked. The respiratory block triggered an immediate gene-expression response diagnostic of TOR inhibition, which was muted by arginine supplementation or without the AMPK-activating kinase Ssp1. The TOR-controlled proteins featured biased composition of amino acids reflecting their shortage after respiratory inhibition. We conclude that respiration supports rapid proliferation in fermenting cells of fission yeast by boosting the supply of Krebs-cycle derived amino acids.

## Introduction

Glucose oxidation in eukaryotic microbes and mammals depends on three core metabolic pathways, glycolysis, the pentose phosphate pathway, and the Krebs cycle. These catabolic processes yield energy in the form of ATP, reducing equivalents in the form of NAD(P)H and FADH, as well as a series of intermediates required for anabolic (biosynthetic) metabolism. The flux distributions between all three pathways are dynamically adjusted by the cell to achieve an optimal level of growth and resilience given a specific lifestyle, stress condition, or ecological niche [1–5]. These metabolic adaptations are of fundamental importance for cellular physiology as they determine their tolerance for changing conditions and stress [6]. Moreover, metabolic flux distributions play a fundamental role in biotechnology. Research has put particular emphasis to explain the balance between respiration and fermentation, i.e. the amount of glucose oxidation achieved via anaerobic metabolism (glycolysis and pentose phosphate pathways without oxidative phosphorylation) versus oxidative metabolism, in which all three pathways are active and feed electrons into the oxygen-consuming respiratory chain [7].

Particularly in eukaryotes, rapidly proliferating cells often use anaerobic metabolism even in the presence of oxygen, called aerobic glycolysis, albeit only full glucose oxidation through the respiratory chain maxes out the redox potential stored in glucose and yields the maximum number of ATP molecules. Growth by aerobic glycolysis is often referred to as Warburg effect in cancer cells [8–10]. Related to aerobic glycolysis is the Crabtree effect in yeast cells, reflecting an active suppression of carbohydrate oxidation in the respiratory chain as long as glucose is available [11,12]. Conversely, differentiated mammalian cells or yeast cells grown in nutrient-poor media shift their energy metabolism towards respiration [8]. Thus the levels of respiration and fermentation are re-balanced in response to physiological and environmental conditions. ATP need alone is insufficient to predict these metabolic programmes. Better predictability of metabolic states has been achieved with resource allocation models [13], accounting for the cost of synthesizing the enzymes [14], changes in the surface-to-volume ratio [15], and thermodynamic constraints such as metabolic energy dissipation [16]. These models arrive at the common conclusion that glycolysis is the more efficient pathway, and that additional glucose oxidation through the respiratory chain might provide only additional benefit when carbon availability is limited, competition is high, or under conditions where the excretion of small sugars has a negative impact, i.e. when cells require ATP without the need to grow.

Besides providing electron donors for generating ATP, the mitochondrial Krebs cycle is crucial for the synthesis of amino acids and other small biomolecules needed for new cell mass [17]. Some respiration is required to ensure Krebs cycle activity for the efficient proliferation of human cancer cells growing by aerobic glycolysis [18]. Upon blocking the respiratory chain by deletion of the mitochondrial genome, cultured mammalian cells need supplementation of uridine and pyruvate for proliferation [19]. Uridine synthesis depends on a functional electron transport chain [20], and pyruvate can rescue defects in respiration-mediated aspartate synthesis [21,22]. Thus, metabolic products from the Krebs cycle can become limiting for mammalian cell proliferation.

Some single-celled organisms can successfully proliferate in the absence of oxidative phosphorylation. Budding yeast (*Saccharomyces cerevisiae*) is a “petite-positive” yeast, which can grow in anaerobic conditions in the absence of respiration [23,24]. Like uridine and pyruvate supplemented mammalian cells, these yeasts can grow even without a mitochondrial genome [25,26]. *S. cerevisiae* has evolved adaptations that loosen its dependence on oxidative phosphorylation. An example is the cytoplasmic location of dihydroorotate dehydrogenase, an enzyme that uncouples uridine synthesis from mitochondrial respiration [27,28]. Other examples are the synthesis of glutamic acid and aspartate which normally requires the Krebs cycle metabolites alpha-ketoglutarate and oxaloacetate, respectively. When growing under fermentative conditions, the carbon flux in the Krebs cycle splits to provide alpha-ketoglutarate in the oxidative branch and fumarate in the reductive branch, while the flux between alpha-ketoglutarate and oxaloacetate is inhibited [29,30]. This strategy allows elimination of succinate dehydrogenase activity and helps to maintain redox homeostasis when the electron transport chain is inhibited [31]. Moreover, when respiration is blocked, *S. cerevisiae* uses the retrograde pathway to activate the glyoxylate cycle that helps to replenish Krebs cycle metabolites [32,33].

However, such adaptations are the exception: most yeast species, including fission yeast (*Schizosaccharomyces pombe*), are petite-negative and do not cope well under anaerobic conditions. The *S. pombe* laboratory strain is part of a group of natural isolates that show higher respiration during fermentative growth, which is caused by a naturally occurring mutation in pyruvate kinase that limits glycolytic flux [34]. We have also reported that respiration is required for rapid cell proliferation during fermentation [35]. Here we define the respiratory functions in fission yeast that support rapid proliferation. We show that respiration during fermentative growth in *S. pombe* is required for the synthesis of arginine and amino acids derived from alpha-ketoglutarate. The synthesis of these amino acids becomes limiting for cell proliferation if the respiratory chain is blocked. We further show that blocking respiration under these conditions leads to the transient inhibition of target of rapamycin (TOR) and longer-term amino acid auxotrophy. Respiration in *S. pombe* cells growing with glucose hence compensates for insufficient amino-acid metabolism.

## Results

### Arginine supplementation is necessary and sufficient for rapid cell growth without respiration

We have shown that antimycin A, an inhibitor of the respiratory chain, slows down the growth of *S. pombe* cells in defined minimal medium, but much less so in rich yeast extract medium with the same glucose concentration [35] (Fig. S1A). This finding suggests that the minimal medium lacks a supplement that becomes limiting when the respiratory chain is inhibited. Indeed, supplementation with soluble amino acids largely rescued this slow growth phenotype (Fig. 1A, Fig. S1A). This result indicates that respiratory inhibition leads to amino-acid auxotrophy that limits cell growth. To test whether any single amino acid is sufficient to rescue the slow growth during fermentation, we grew cells in minimal medium with antimycin A and supplemented with 18 individual amino acids; as a control, we used medium supplemented with the mix of all amino acids or without any supplementation. Notably, supplementation with arginine (R) alone could promote growth nearly as much as the mix of all amino acids (Fig. 1A). This effect was dependent on the dose of arginine (Fig. S1B). Thus arginine, but no other single amino acid, is sufficient to promote rapid cell growth when the respiratory chain is inhibited.

**Figure 1.**
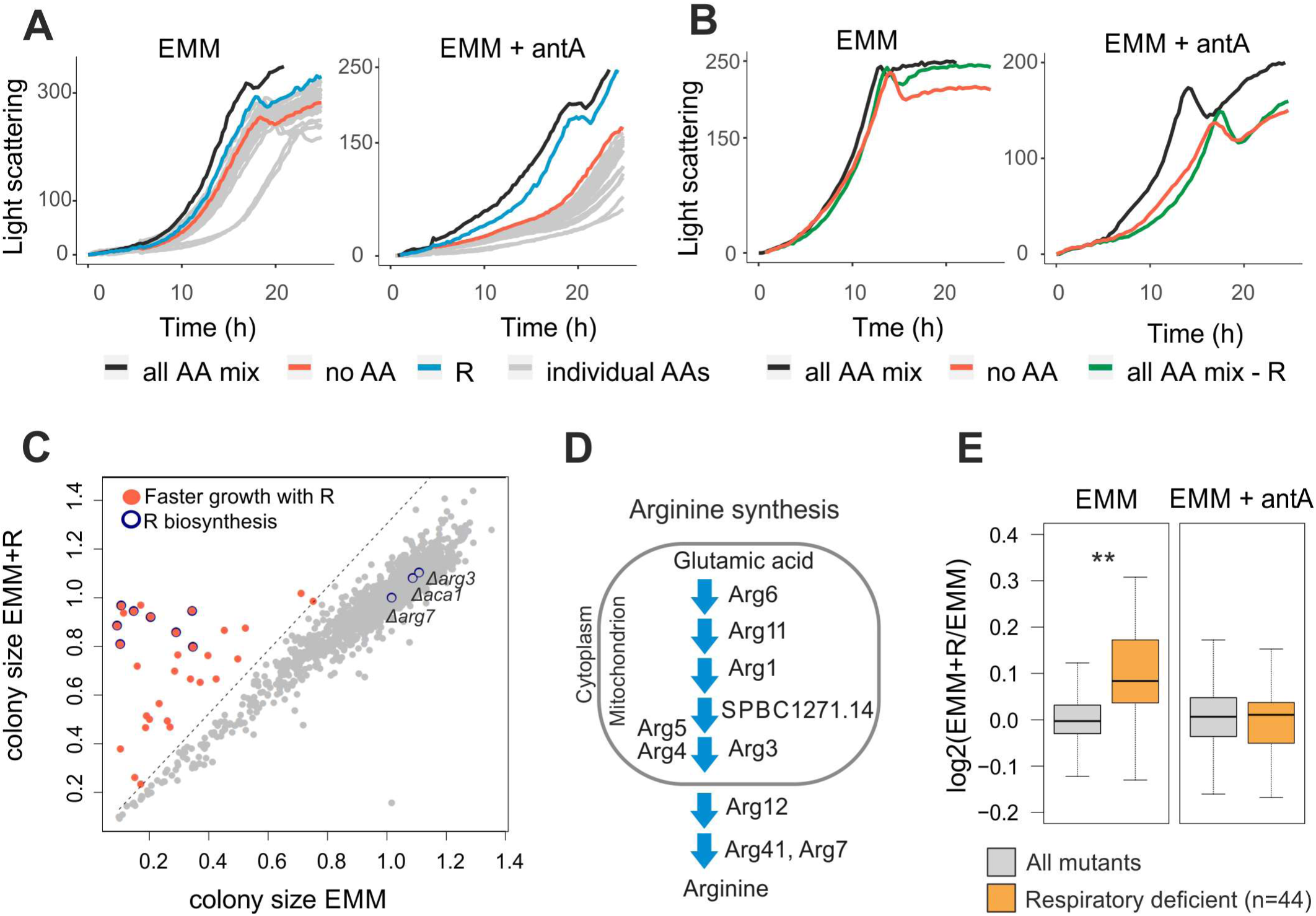
Arginine enhances growth of cells with inhibited respiratory chain. **(A)** Growth curves of *S. pombe* cultures in minimal medium without (EMM) or with antimycin A (EMM + antA). Cultures were supplemented with either the complete amino-acid mix (all AA mix, black), without any amino acids (no AA, red), with arginine only (R, blue), or with individual amino acids other than arginine (individual AAs, grey). **(B)** Growth curves as in (A) for cultures supplemented with complete amino-mix (black), without any amino acids (red), or amino-acid mix without arginine (green). **(C)** Scatter plot of normalised colony sizes of cells grown on EMM against normalised colony sizes of cells grown on EMM supplemented with arginine (EMM+R) (Table S1). Mutants that passed our filtering (Methods) and whose colony-size ratios were 30% bigger when supplemented with arginine are shown in red; mutants encoding known genes of the arginine biosynthesis pathway are shown with blue circles. **(D)** Scheme of arginine biosynthesis pathway, with protein localisation based on PomBase annotations [59]. **(E)** Box plot of colony size ratios of mutants grown with *vs* without arginine for all mutants (grey) and respiratory deficient mutants (orange, based on previous screens [37] (Welch’s t-test *pval* < 0.002). The same analysis was performed for mutants grown on EMM (left) or on EMM supplemented with antimycin A (right) (Table S2).

To test whether arginine is necessary for rapid cell growth with inhibited respiratory chain, we grew cells with all amino acids except arginine. While supplementation of the mix of all amino acids allowed rapid growth, omission of arginine from the mix led to slower growth similar to cells not supplemented with any amino acids (Fig. 1B). We conclude that cell growth in minimal medium is reduced upon blocking the respiratory chain, and arginine supplementation is both sufficient and necessary to increase growth of respiring cells to near normal levels.

These results suggest that respiration is important for the synthesis of arginine. We therefore expected that respiration-defective mutants are auxotroph for arginine. To test this hypothesis, we screened for arginine-auxotroph mutants by growing the ∼3500 strains of the haploid *S. pombe* gene-deletion library in the presence or absence of arginine. This screen identified 31 mutants that grew substantially better when supplemented with arginine (Fig. 1C; Table S1). As expected, these arginine-auxotroph mutants included those with impaired arginine biosynthesis. Out of 11 genes associated with the Gene Ontology (GO) category ‘arginine biosynthesis’ (GO:0006526), 8 were identified in our screen (Fig. 1C,D). Of the remaining three, *aca1* does not have a confirmed function in arginine metabolism, *arg7* is one of two arginosuccinate lyases (with deletion of the other one, *arg41*, showing arginine auxotrophy), and *arg3* is wrongly annotated in the deletion library, because an independently prepared deletion strain showed arginine auxotrophy (Fig. S4A). Notably, most steps of the arginine biosynthesis pathway occur in mitochondria (Fig. 1D).

The screen identified 23 additional genes not previously associated with arginine metabolism. Like the arginine-biosynthesis genes, these 23 genes were enriched for those encoding mitochondrial proteins (13 of 23; *p* <1E-5); some of these 13 genes encode mitochondrial metabolic enzymes (e.g., Lat1, Maa1, Idh1, Pos5, Atd1) whose deletion could impact arginine synthesis by affecting the Krebs cycle and glutamic acid production (Fig. S2A), while other genes encode respiratory functions like subunits of the mitochondrial ATP-ase or proteins involved in mitochondrial transcription (Ppr7) or translation (Mpa1). Our screen also identified a mutant of mitochondrial thioredoxin (Trx2) which is known to be rescued by arginine [36]. Threshold-free analysis of the screen results further revealed significant enrichment of genes encoding mitochondrial proteins among the strains that grew better with arginine (Fig. S2B). Thus, arginine auxotrophy can be caused not only by mutants directly affecting arginine synthesis but also by mutants affecting different mitochondrial functions.

Given that arginine was not essential for cell growth after blocking the respiratory chain (Fig. 1A), some respiratory mutants may show only subtle growth defects in the absence of arginine. Using data from our previous screen [37], we checked how a group of 44 respiratory deficient mutants grew in the screen reported here. Overall, the growth of respiratory mutants was significantly enhanced by arginine (Fig. 1E). As a control, we screened all mutants in the presence or absence of arginine but with antimycin A to block the respiratory chain in all strains. Under this condition, we did not detect any growth differences between respiratory and other mutants, indicating that growth inhibition without arginine was indeed caused by differences in respiration (Fig. 1E; Table S2). We also checked for effects of arginine on the growth of respiratory mutants in liquid medium: out of 44 respiratory mutants, 20 exhibited a growth phenotype in minimal medium (either slow growth or lower final biomass), while arginine supplementation attenuated the growth phenotype for 15 of these mutants. (Fig. S3). We conclude that many respiratory mutants become dependent on arginine, indicating that mitochondrial respiration is important for arginine biosynthesis.

### Block of respiratory chain leads to changing cellular amino-acid levels

To further analyse the link between respiration and arginine metabolism, we measured the levels of free amino acids in cells treated with antimycin A. To this end, we used liquid chromatography selective reaction monitoring [38] to determine intracellular concentrations of 16 amino acids (Table S3). Indeed, cells with blocked respiratory chain showed about 9-fold reduced levels of arginine (Fig. 2A). In addition, these cells showed reduced levels of ornithine (Orn), a metabolite of the arginine synthesis pathway, and of glutamic acid (E), a substrate for arginine biosynthesis [39] (Fig. 2A). In contrast, levels of alanine (A) were elevated about 2.5-fold (Fig. 2A). Interestingly, plants exposed to hypoxic conditions also accumulate alanine, which might serve as carbon/nitrogen reservoir [40].

**Figure 2.**
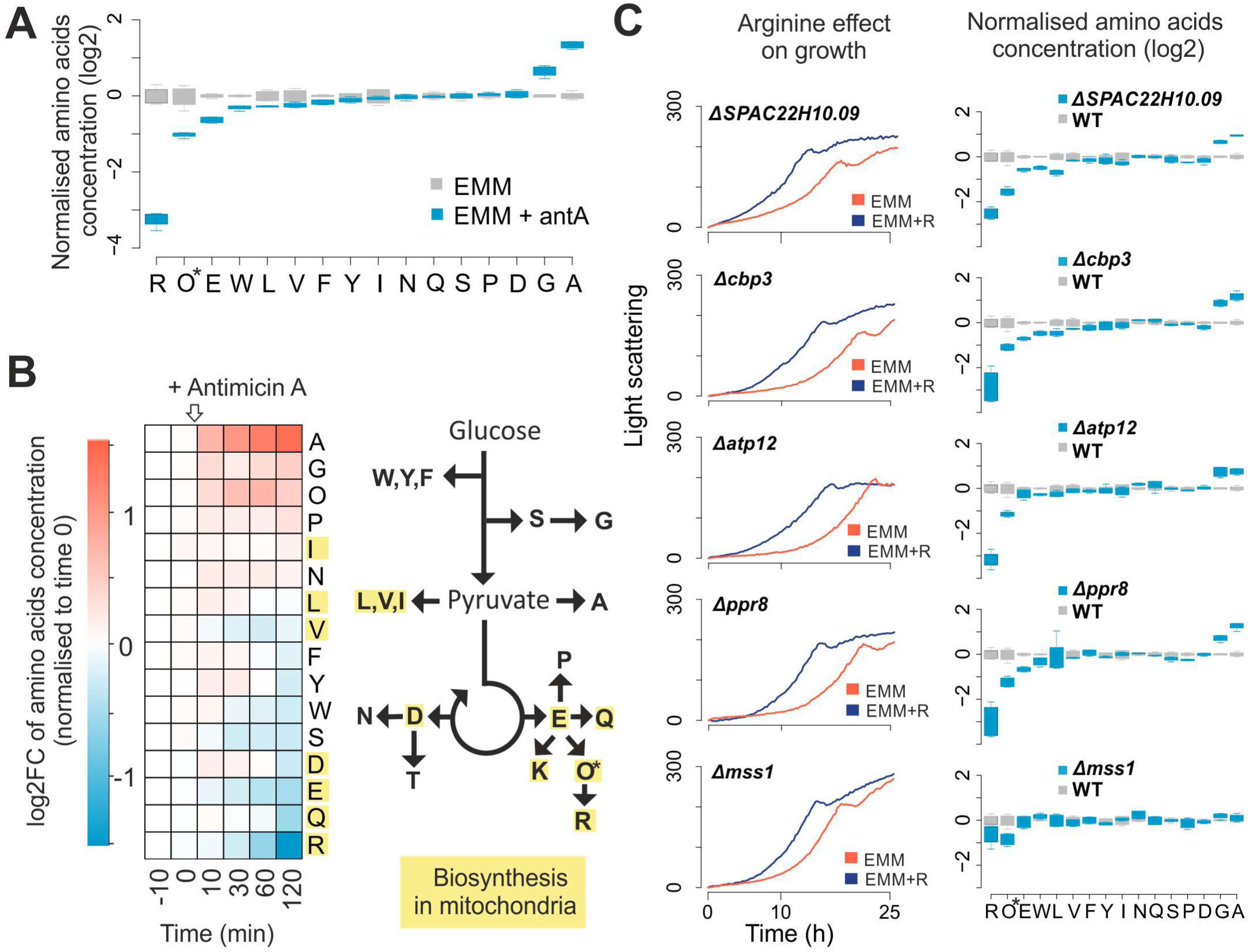
Respiratory inhibition alters levels of cellular amino acids. **(A)** Normalised concentration values of 16 amino acids in cells grown in EMM without (grey) or with (blue) antimycin A. For each condition, four independent samples were analysed. Concentrations are presented relative to the mean value for the given amino acid in EMM medium. *Ornithine (Orn) is abbreviated as O. **(B)** Amino-acid concentrations before and after addition of antimycin A to cells exponentially growing in EMM. Left: heat map showing concentration changes (log2 fold change) of each measured amino acid during the time course relative to time point 0 (before antimycin A addition). Right: simplified scheme of amino-acid biosynthetic pathways. Amino acids for which any biosynthetic enzyme has an annotated mitochondrial location are highlighted in yellow. *Ornithine (Orn) is abbreviated as O. **(C)** Left panels: cell growth in EMM without (red) or with (blue) arginine supplementation for five selected respiratory mutants as indicated. Right panels: normalised intracellular amino-acid concentrations as in (A) for the same 5 respiratory mutants (blue) and wild-type control cells (grey). For each mutant, four independent samples were analysed. *Ornithine (Orn) is abbreviated as O.

We also measured the dynamic changes in free intracellular amino acids within 2 hours after exposing cells to antimycin A. Consistent with cells grown in steady-state, inhibition of the respiratory chain led to increasing levels of alanine and decreasing levels of arginine (Fig. 2B). Notably, several other amino acids decreased, mostly derivatives of Krebs-cycle metabolites such as glutamic acid and glutamine. To confirm that the altered amino-acid levels are not an artefact of antimycin A treatment, we measured amino-acid concentrations in five respiratory mutants whose growth was improved by arginine (Fig. S3). In these mutants, except *mss1*, the patterns of amino-acid changes were similar to those of cells grown with antimycin A (Fig. 2C). Most notably, the levels of arginine were lowered while the levels of alanine increased (Fig. 2C). These results corroborate that the slow growth of respiratory mutants in minimal medium is caused at least in part by limiting intracellular levels of arginine. Together, these findings show that respiration is important for the synthesis of arginine and some other amino acids that are dependent on the Krebs cycle.

### Amino acids derived from arginine contribute to rapid cell growth with blocked respiratory chain

The changes in intracellular amino-acid concentrations pointed to additional effects from arginine supplementation: ornithine levels transiently increased, peaking 60 minutes after blocking the respiratory chain (Fig. 2B), while they decreased after an extended respiratory block (Fig. 2A). This result suggested that arginine synthesis may be blocked at the transition from ornithine to citrulline or that cellular arginine is catabolised to ornithine. The first possibility is consistent with a report showing that absence of mitochondrial thioredoxin results in lowered Arg3 which catalyses the ornithine-to-citrulline transition [36]. In our hands, however, neither the localisation of Arg3 nor its protein or RNA levels did change after antimycin A treatment or in respiratory mutants (Fig. S4). Arginine can be catabolised to other amino acids, a conversion that is highly efficient in *S. pombe* with arginine being rapidly catabolised to proline, glutamate, glutamic acid and lysine [41]. Accordingly, cells supplemented with arginine showed not only higher levels of arginine but also of amino acids produced by arginine catabolism: ornithine, proline and glutamic acid (Fig. 3A). Thus, upon blocking of respiration and associated arginine synthesis, arginine catabolism may transiently increase the levels of ornithine (Fig. 2B), which in turn is converted to other amino acids (Fig. 3B). These findings raised the possibility that metabolites derived from arginine, besides the supplemented arginine itself, contribute to the rapid cell growth when the respiratory chain is inhibited (Fig. 1).

**Figure 3.**
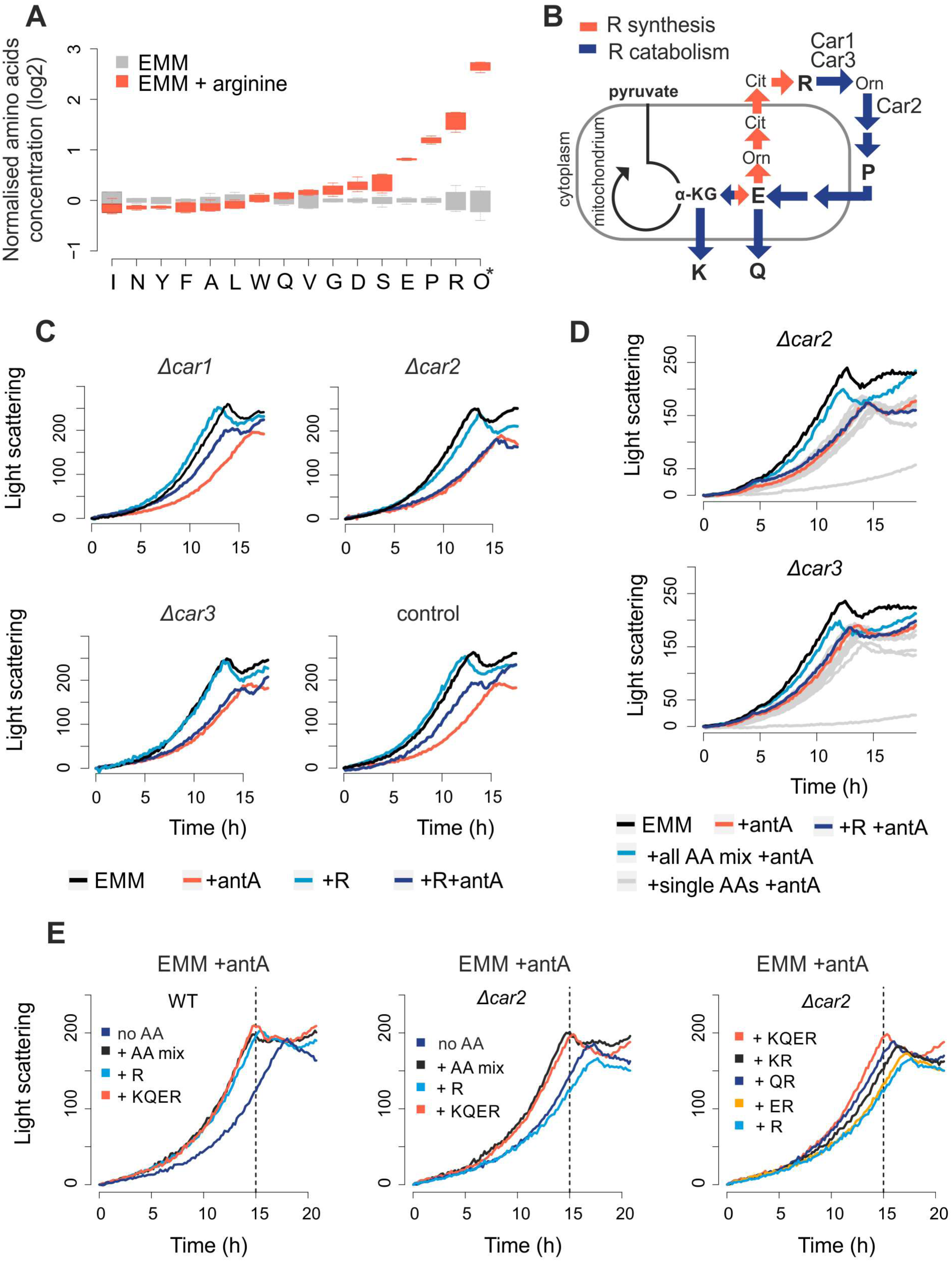
Arginine catabolism contributes to growth of cells with blocked respiratory chain. **(A)** Normalised amino-acid concentrations in cells grown in EMM without (grey) or with arginine (red). For each condition, four independent samples were analysed. Concentrations are presented relative to the mean value for the given amino acid in EMM medium. *Ornithine (Orn) is abbreviated as O. **(B)** Schematic relationship between alpha-ketoglutarate (α-KG) and arginine (R) synthesis (red arrows) and catabolism (blue arrows). Arginine is used to synthesize proline (P), glutamic acid (E), glutamine (Q) and lysine (K). The enzymatic roles of Car1, Car2 and Car3 are indicated. **(C)** Growth curves of strains deleted for *car1, car2* or *car3*, and a wild-type control. Strains were grown in EMM (black), EMM with antimycin A (red), EMM with arginine (light blue) or EMM with both arginine and antimycin A (dark blue) as indicated. **(D)** Growth curves of strains deleted for *car2* (top) or *car3* (bottom). Strains were grown in EMM (black), EMM with antimycin A (red), EMM with arginine and antimycin A (dark blue), EMM with mix of all amino acids and antimycin A (bright blue), and EMM with antimycin A and each individual amino acid from the mix (grey). **(E)** Growth curves of strains deleted for *car2* or *car3*, and a wild-type control (WT). Strains were grown in EMM media with antimycin A and with either a mix of all amino acids (+AA mix) or different combinations of lysine (K), glutamine (Q), glutamic acid (E), or arginine (R) as indicated.

To investigate the role of arginine catabolism after blocking the respiratory chain, we constructed three mutants deleted for the genes encoding the arginine-catabolism enzymes Car1, Car2 and Car3. Like the wild-type cells, all three mutants showed reduced growth upon antimycin A treatment (Fig. 3C). Arginine supplementation did enhance growth in wild-type and *car1* deletion cells, but not in *car2* and *car3* deletion cells (Fig. 3C). This difference between the mutants may reflect that Car1 only plays a negligible role in arginine catabolism in standard conditions [41]. Our results show that arginine catabolism is required for rapid proliferation of arginine-supplemented cells with inhibited respiratory chain.

We checked what arginine-derived amino acids are required for rapid cell proliferation when blocking the respiratory chain. While no single amino acid could increase the growth of *car2* or *car3* mutants, a mix of all amino acids did increase the growth of both mutants (Fig. 3D). To establish which amino-acid combinations are necessary to increase growth without respiration, we used the *car2* mutant which exhibited a stronger growth phenotype. Arginine combined with its catabolic products, glutamine, glutamic acid and lysine, led to full restoration of cell growth (Fig. 3E). Of the three arginine-derived amino acids, glutamine and lysine led to stronger growth increases than glutamic acid. Taken together, these results show that respiratory chain inhibition does not result only in inhibition of arginine synthesis, but also in a shortage of glutamine, lysine and glutamic acid. All these amino acids are derivatives of alpha-ketoglutarate produced by the Krebs cycle, indicating that the reduced growth is caused by inhibition of the Krebs cycle. Thus, inhibition of the electron transport chain through antimycin A also triggers and inhibition of the Krebs cycle. This contrasts the situation in budding yeast where the Krebs cycle can be uncoupled from the activity of the electron transport chain [42].

### Transcriptome regulation in response to respiratory chain block and arginine

Inhibition of the respiratory chain leads to a retrograde response, including the downregulation of genes for subunits of the electron transport chain [35,43]. Given the impact of respiratory chain inhibition on amino-acid homeostasis (Fig. 2), we used microarrays to investigate transcriptome changes within 2 hours after antimycin A addition in the presence or absence of arginine. Addition of antimycin A inhibited the expression of genes functioning in the electron transport chain regardless of arginine supplementation (Fig. S5). Moreover, antimycin A resulted in an immediate, strong Core Environmental Stress Response (CESR) [44], regardless of arginine supplementation (Fig. 4A; Table S4).

**Figure 4.**
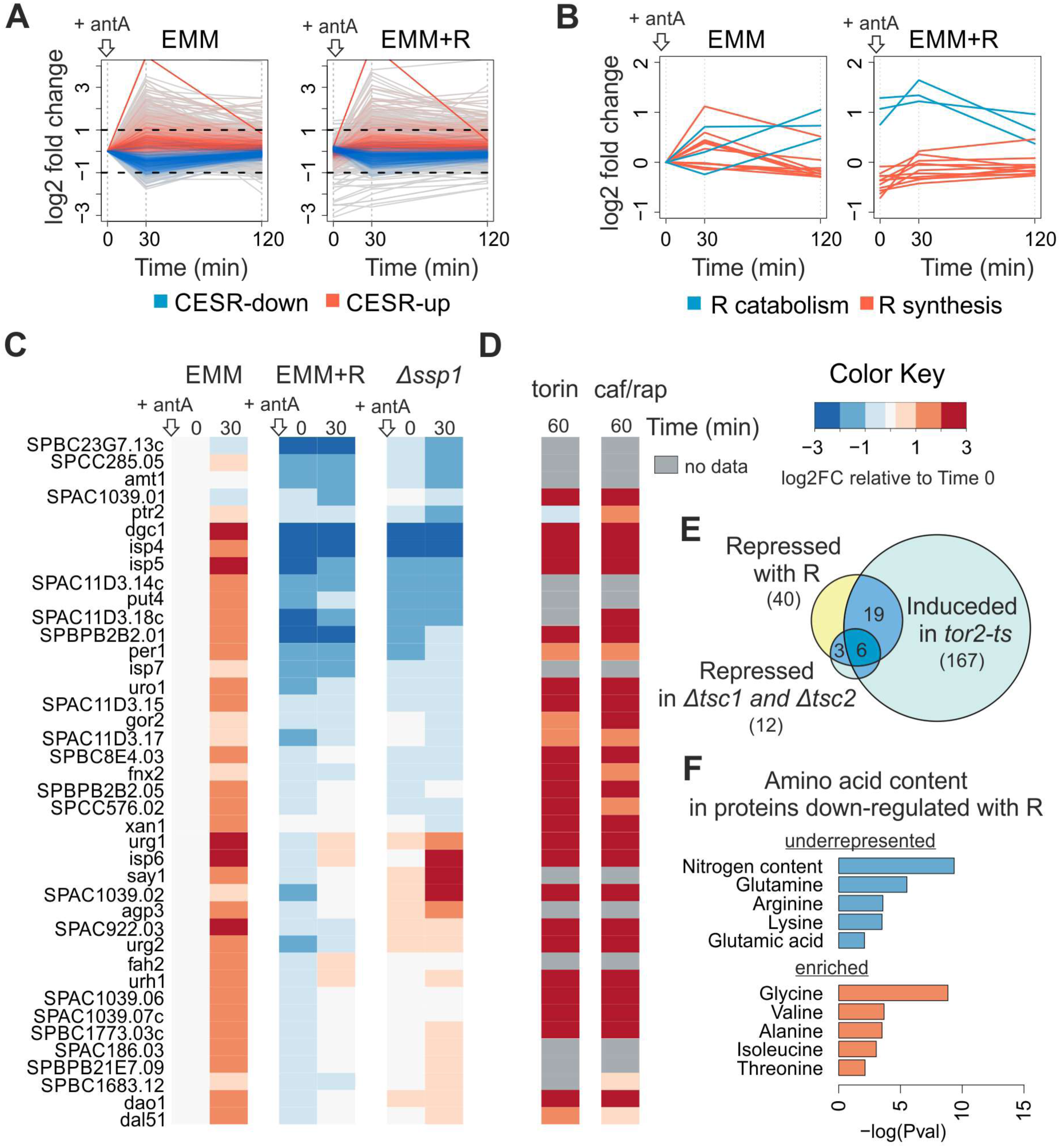
Transcriptome regulation in response to respiratory inhibition and arginine supplementation. **(A)** Changes in transcript abundance 30 and 120 min after addition of antimycin A to exponentially growing cells without (EMM) or with arginine (EMM+R). Transcripts annotated as up- or down-regulated Core Environmental Stress Response (CESR) are indicated in red and blue, respectively. All data is normalised to EMM time point 0. **(B)** Heat map of expression changes for transcripts repressed over 2-fold at 30 min in cells supplemented with arginine (EMM +R) relative to cells without supplementation (EMM). Values for timepoints 0 and 30 min after antimycin A addition are shown for cells grown in EMM, EMM with arginine and *ssp1* deletion mutant grown in EMM. The data are normalised to EMM timepoint 0. **(C)** Heat map of expression changes of the same genes as in (B) after one hour of inhibiting TOR kinase with torin1 or caffeine/rapamycin treatment relative to cell before treatment; cells were grown in rich media - YES (data from [45]) **(D)** Overlaps between genes repressed with arginine supplementation and genes induced in *tor2* thermo-sensitive mutant (p <E-27) [48], or repressed in strains deleted for the TOR inhibitors *tsc1* and *tsc2* (p <E-16) [47]. **(E)** Enriched and underrepresented amino acids in proteins encoded by 40 genes that are down-regulated in media with arginine. Analysis performed using online tool AnGeLi [60].

Interestingly, after addition of antimycin A, a group of 40 genes were less induced or even inhibited in the presence of arginine (Fig. 4B; Fig. S5B). Most of these genes are known to be induced after inhibition of TOR signalling (Fig. 4C; Fig. S5C) [45]. Accordingly, this group was enriched for genes that are repressed or induced, respectively, upon genetic activation or inhibition of TOR kinase signalling (Fig. 4D) [46–48]. We therefore conclude that blocking the respiratory chain not only results in a stress response but also leads to a transient TOR inhibition and up-regulation of genes that are normally inhibited by TOR.

Arginine can activate TOR signalling [49,50]; thus the lower expression of the 40 genes in arginine-supplemented cells may reflect that arginine antagonized the TOR inhibition triggered by the respiratory chain block. This effect of arginine could potentially explain its pro-growth effect when the respiratory chain is blocked. However, TOR-inhibited transcripts genes were up-regulated within 30 minutes (Fig. 4B), while arginine concentration substantially decreased only 2 hours after antimycin A treatment (Fig. 2B). It is therefore more likely that arginine enhances growth of respiration-inhibited cells simply by replenishing the limiting amino acids. This conclusion is supported by our finding that supplementation with other amino acids that activate TOR, like lysine or glutamine, did not promote cell proliferation (Fig. 1A).

Given that the rapid, transient TOR inhibition is unlikely to result from the lower arginine levels in response to a respiratory inhibition, it may be triggered by a different signal. We hypothesized that the AMP-activated protein kinase (AMPK) could mediate such an earlier signal. Consistent with this idea, antimycin A-treated cells without the AMPK-activating kinase Ssp1 did not up-regulate most of the 40 genes that were repressed by arginine (Fig. 4C). This result suggests that TOR is indeed inhibited by AMPK kinase upon blocking respiration. Consistently, regulation of TOR via AMPK in response to environmental perturbations has been reported in fission yeast [50,51], and lowering the Krebs cycle flux leads to AMPK-mediated TOR inhibition in budding yeast [42]. After blocking respiration, the cells may bounce back from the transient TOR inhibition, through a retrograde response and redirection of energy metabolism to fermentation, until the levels of arginine and derived amino acids become limiting for rapid proliferation.

The protein products of the 40 TOR-regulated genes that are repressed by arginine supplementation (and induced by antimycin A) featured lower than average contents of nitrogen, arginine, lysine, glutamine and glutamic acid (Fig. 4E). Strikingly, these amino acids are all derived from alpha-ketoglutarate which becomes scarce without sufficient flux in the Krebs cycle (Fig. 2B). This finding illustrates the relationship between available resources and cell regulation. So a block in the respiratory chain results in shortage of amino acids derived from alpha-ketoglutarate, and proteins that are induced to deal with this shortage, including amino-acid transporters; Fig. S5C, feature low levels of these limiting amino acids. On the other hand, these induced proteins featured higher than average contents of alanine, isoleucine, valine, glycine and threonine, consistent with the intracellular increase in alanine and glycine in the absence of respiration (Fig. 2B). These findings indicate that the gene expression response to altered levels of arginine or respiration involves proteins whose content is adapted to intracellular amino-acid availability in these conditions.

## Conclusion

We demonstrate here that inhibiting the respiratory chain generates a fitness cost for *S. pombe* cells grown in fermentative conditions. Respiration contributes to rapid proliferation during aerobic fermentation by supplying amino-acids derived from alpha-ketoglutarate, a Krebs cycle metabolite. Respiratory inhibition therefore leads to a shortage of arginine, glutamic acid, glutamine and lysine. This shortage can be largely alleviated by supplementation with arginine which is efficiently converted to other alpha-ketoglutarate derivatives. Our transcriptome analysis shows not only a retrograde response but also a transient, rapid response to stress and TOR inhibition after blocking the respiratory chain. An intact AMPK pathway is required for this TOR inhibition. Budding yeast features a similar response to a Krebs-cycle block, but not to a respiration block [42].

Some fission yeast isolates, including the standard laboratory strain, possess a mutation in pyruvate kinase, a key glycolytic enzyme [34]. This feature limits glycolytic flux and causes increased respiration during fermentative growth [34]. It is possible that the shortage of amino acids is exacerbated by this mutation. *S. pombe* thus provides a unique model to study how increased respiration compensates for lower glycolytic flux. Rapid proliferation is usually supported by fermentative metabolism even in the presence of oxygen. However, aerobic glycolysis does not replace respiration, and it becomes clear that both processes occur in parallel to support the metabolic needs of cells. We show here that respiration in *S. pombe* is required to supply amino-acid derivatives of the Krebs cycle. Arginine is necessary and sufficient to supress the growth inhibition triggered by blocking the respiratory chain. In human cells, respiration limits proliferation due to its contribution to aspartate synthesis [21,22]. Intriguingly, although arginine is essential in humans, several reports indicate positive effects of arginine supplementation on patients with mitochondrial dysfunction [52].

## Materials and Methods

### Yeast strains and growth media

Strains used in this study are listed in Table S5. For colony size measurements, the auxotroph Bioneer library v5.0 was used [53]. Respiratory mutants were isolated from prototroph derivative of auxotroph collection [37]. Cells were grown in rich Yeast Extract with Supplements (YES) medium with 3% glucose or defined Edinburgh Minimal Medium (EMM) with 2% glucose (Formedium PMD12CFG). Antimycin A was used where indicated at 0.15 μg/ml concentration. Amino acids were used at 0.2 g/L concentration each unless indicated otherwise.

### Growth recording and analysis

Growth curves of cells growing in liquid media were recorded using BioLector (m2P-labs) microfermentor. Data were normalised to time zero and visualised using custom R scripts. For colony size measurement, deletion library was arrayed on solid agar EMM media plates in in 384 colony/plate format using a RoToR HDA robot (Singer Instruments). After two days of growth plates were scanned and raw colony sizes extracted using gitter R package [54]. Colony sizes were normalised using an in-house R package (HDAr package - unpublished). Plates were supplemented with arginine or antimycin A. Additionally plates were supplemented with adenine, uracil and leucine. Experiment with arginine supplementation was performed in four repeats (two parallel repeats in two different batches). Experiment with arginine supplementation in presence of antimycin A was performed in two parallel repeats. All normalised colony sizes from aforementioned experiments are provided in Tables S1 and S2. To identify strains growing better on arginine, we extracted colony size information from all EMM and EMM+R plates and removed data for most variable colonies for each condition (coefficient of variation >0.25).

### Amino acids measurement

Intracellular amino acids were quantified as previously described [38,55,56]. In brief, samples were prepared by hot ethanol extraction of mid-log phase cultures. Then, 1.5 ml l of culture were spun down, washed with deionised water (3000 g, 3 mins), and 200 μL of ethanol at 80°C were added. The pellet was resuspended by vortexing and incubated for 2 min at 80°C, followed by vortexting and another 2 min incubation step. The extract was cleared by centrifugation at 3000 g for 5 min, and the supernatant was used for LC-MS analysis without further conditioning. Chromatographic separation was performed on 1 μl of sample using a Waters ACQUITY UPLC BEH amide 1.7 mm 2.1 × 100 mm column at 25 °C. Flow was maintained at 0.9 ml min^−1^, starting with 15% Buffer A (50:50 acetonitrile/water, 10 mM ammonium formate, 0.176% formic acid) and 85% Buffer B (95:5:5 acetonitrile/methanol/water, 10 mM ammonium formate and 0.176% formic acid) for 7 min, followed by a 1.85-min ramping to 5% B, and keeping constant for 0.05 min before returning to the equilibrating conditions of 85% B. Quantification was carried out by multiple reaction monitoring (MRM) on a triple quadrupole mass spectrometer (Agilent 6460). Peak areas were converted to concentrations by external calibration with amino acid standard mixtures. Cysteine, methionine, lysine and histidine could not be resolved or quantified reliably and were excluded from further analysis. Data was normalised by median quotient normalisation as described [57].

### Microarray experiments

For the time course analyses, cells were grown to early exponential phase (OD 0.5) in EMM medium with or without 0.2 g/L arginine. Transcriptomes were analysed before (time point 0) and at two time points after adding antimycin A to the culture (30 and 120 minutes). Cells were collected using centrifugation, washed once with ice cold water and snap frozen. RNA from cell pellets was isolated using hot phenol extraction, followed by labelling of the single samples and a pool of all the samples which served as reference [58]. Agilent 4 × 44K custom-made *S. pombe* expression microarrays were used, and hybridizations and subsequent washes were performed according to the manufacturer’s protocols. Microarrays were scanned using a GenePix 4000 B laser scanner, and fluorescence signals were analysed using GenePix Pro software (Axon Instruments). The resulting data were processed using customized R scripts for quality control and normalization and analysed using GeneSpring software. Normalised data from GeneSpring are included in Table S4. All the visualisations were performed based on this data set with use of custom R scripts.

## Acknowledgments

We thank Maria Rodríguez-López for help with microarray analysis, StJohn Townsend for help with colony-size screen analysis, Mimoza Hoti for technical support, and Charalampos Rallis for advice and fruitful discussions. We thank Shajahan Anver, Snezhka Oliferenko, Charalampos Rallis and Maria Rodríguez-López for comments on the manuscript. This work was supported by a Wellcome Trust Senior Investigator Award to J.B. (grant no. 095598/Z/11/Z), by the Foundation for Polish Science FIRST TEAM grant awarded to M.M (POIR.04.04.00-00-4316/17), and by the Francis Crick Institute which receives its core funding from Cancer Research UK (FC001134), the UK Medical Research Council (FC001134) and the Wellcome Trust (FC001134).

## Supplementary figure legends

**Figure S1.**
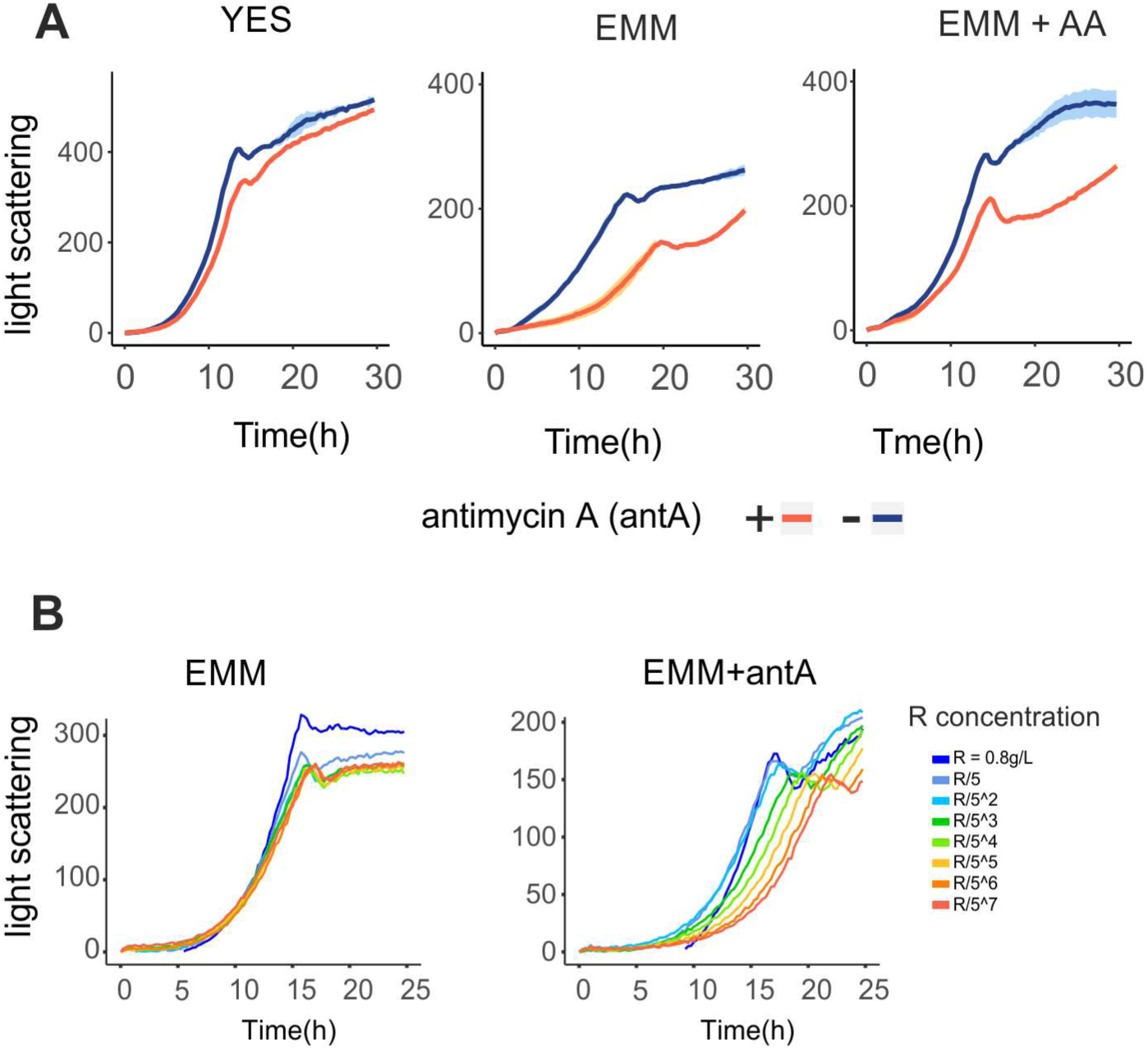
Arginine improves growth of cells with blocked respiration. **(A)** Growth curves of yeast cultures in rich (YES) or defined (EMM) media, and in defined media supplemented with mix of soluble amino acids (EMM + AA). In each experiment media were supplemented or not with antimycin A (blue or red curves respectively). **(B)** Dose dependent effect of arginine. Growth curves of yeast cultures in defined media with or without antimycin A (EMM or EMM + antA), supplemented with different amounts of arginine (as indicated in the legend).

**Figure S2.**
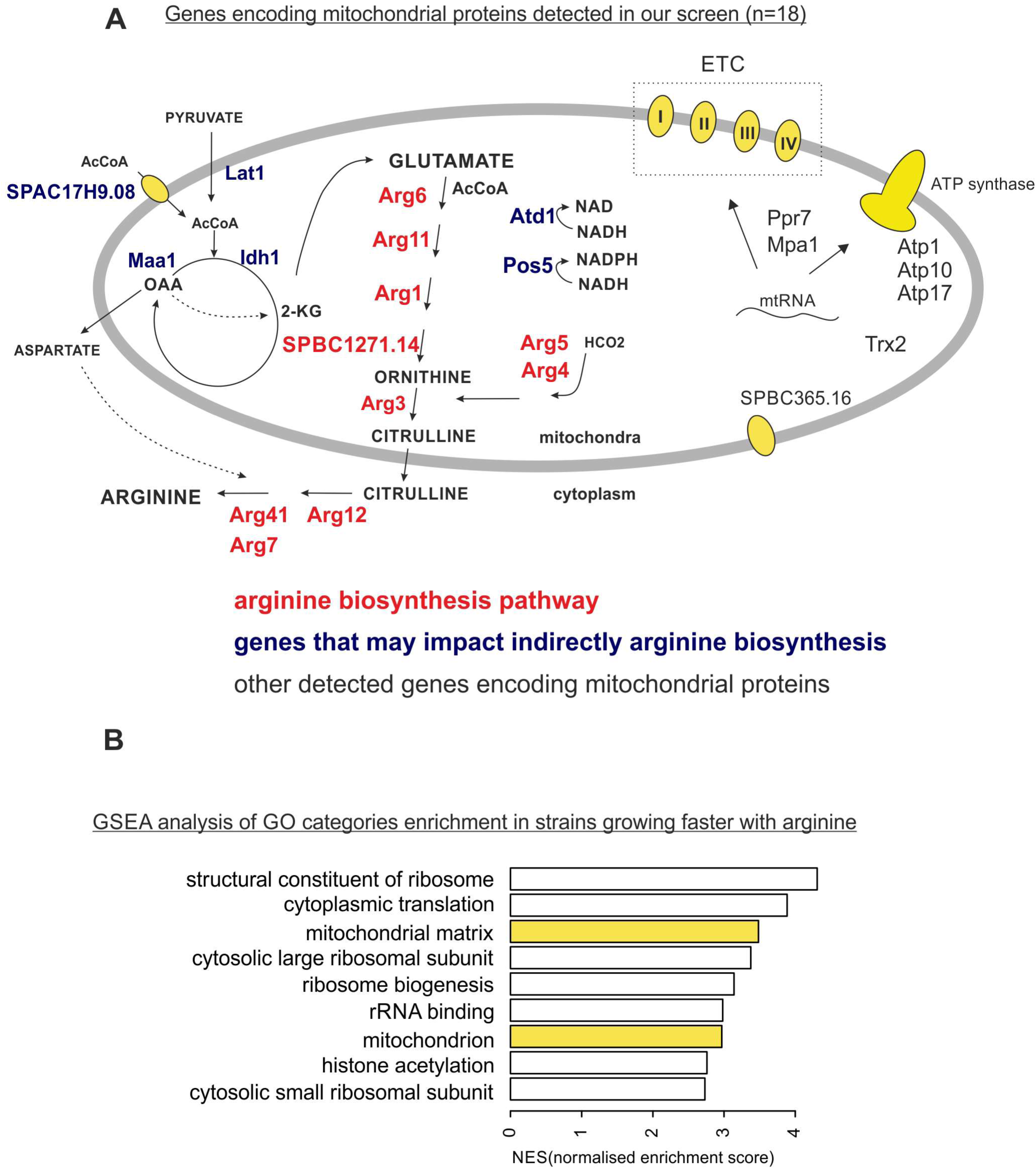
Genes whose deletion resulted in arginine auxotrophy. **(A)** Genes encoding mitochondrial proteins identified in our screen were separated to ones that are part of arginine biosynthesis pathway (red), ones that may indirectly impact both respiration and arginine biosynthesis (blue) and others (black). **(B)** Gene Ontology (GO) enrichments in the result of the screen identified using threshold-free GSEA analysis [61]. (we show categories enriched among colonies growing faster on with arginine).

**Figure S3.**
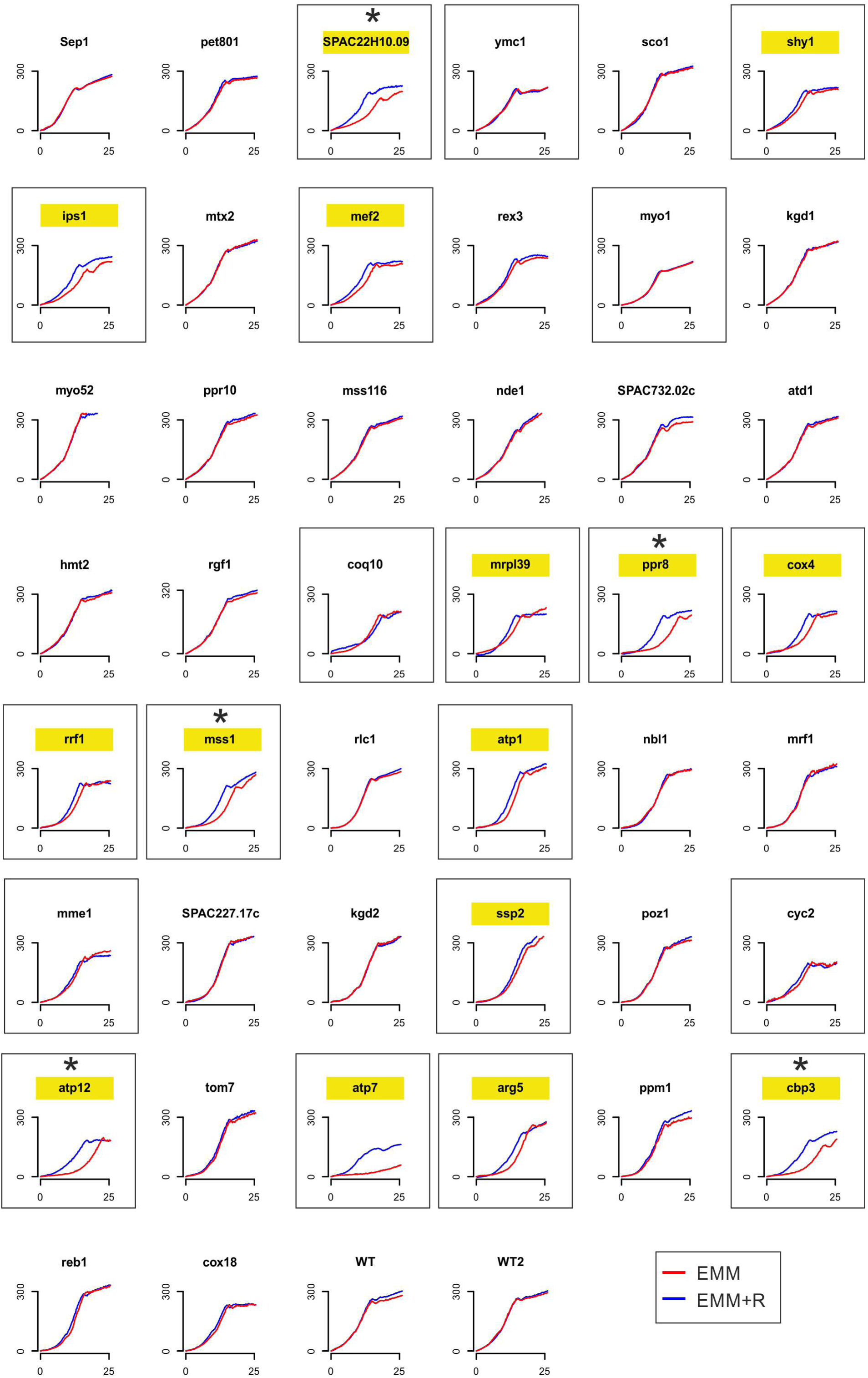
Growth of respiratory mutants can be improved by arginine. Growth curves of 44 respiratory mutants in defined media (EMM) with or without arginine (blue or red respectively). Strains were isolated from prototroph library [37]. In frames are growth curves of strains whose growth was inhibited in minimal media, and highlighted in yellow are strains whose growth was improved by arginine supplementation. Strains that were selected for further experiments are indicated with asterisks.

**Figure S4.**
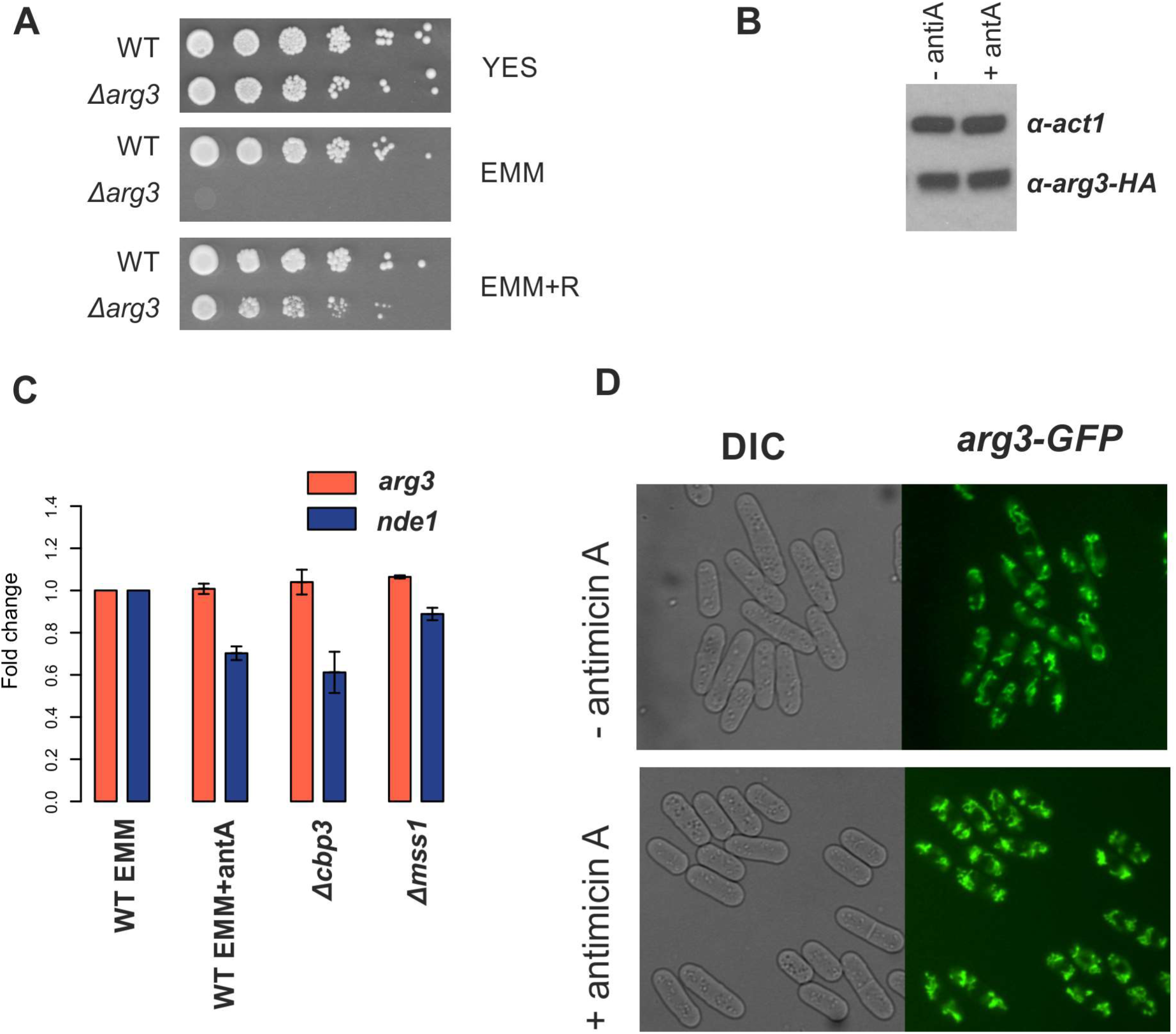
Respiration inhibition does not impact endogenous *arg3* RNA or protein level. (**A**) Strain with *arg3* gene deletion (*Δarg3*) in prototroph background exhibits arginine auxotrophy. Wild type prototroph (WT) and *Δarg3* strains were grown in liquid YES media, washed in EMM and spotted on agar plates with rich (YES) and minimal (EMM) medium, and in minimal medium supplemented with arginine (EMM+R). **(B)** Western blot against Arg3-HA fusion protein. Total proteins were extracted from cells grown exponentially in EMM or EMM with antimycin A. Signal obtained using antibodies against actin (α-act1) serves as loading control. **(C)** Arg3-GFP fusion localises to mitochondria, we detected similar intensity of GFP signal in cells grown in EMM or in EMM with antimycin A. **(D)** Level of *arg3* transcript was measured using qPCR method in cells grown in EMM or EMM with antimycin A. Additionally level of *arg3* transcript was measured in respiratory mutants grown in EMM – *Δcbp1* and *Δmss1*. As a control we measured level of *nde1* transcript encoding equivalent of complex I electron transport chain, that was shown to be repressed in respiratory mutants [35]. Data were normalised to the *cdc2* transcript level.

**Figure S5.**
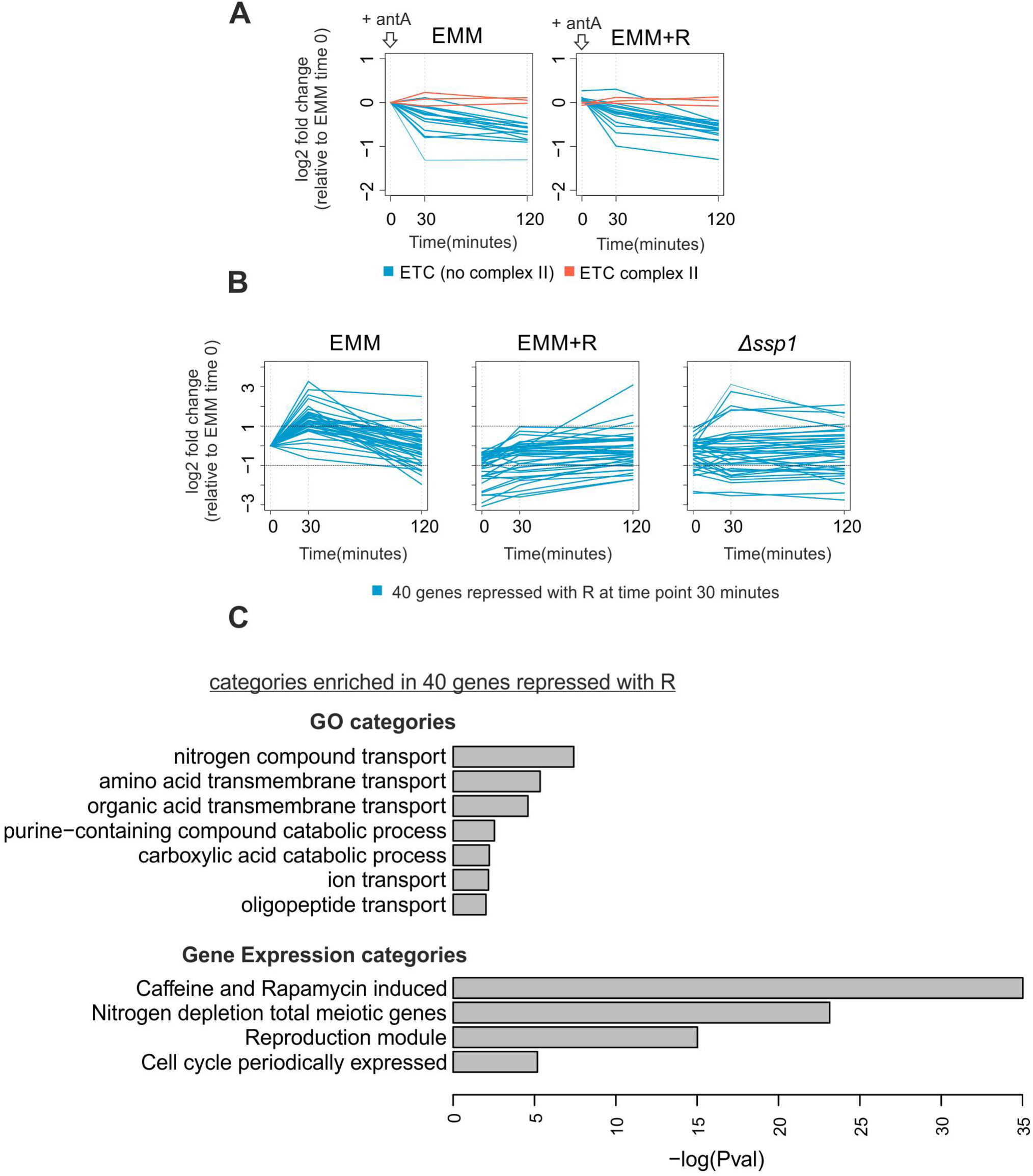
Arginine does not impact repression of genes encoding electron transport chain components. **(A)** Changes of transcripts encoding electron transport chain complex subunits before and 30 and 120 min after antimycin A treatment in minimal media (EMM) and minimal media supplemented with arginine (EMM +R). Subunits of complex II are indicated in red, and other subunits in blue. Values are normalised to EMM time zero. **(B)** Profiles of expression of 40 genes repressed in media with arginine at 30 min after antimycin A treatment. Changes of transcript abundances before and 30 and 120 min after antimycin A treatment are shown for the wild-type cells grown in minimal media with or without arginine and for *ssp1* deletion cells grown in minimal media. Values are normalised to EMM time zero. **(C)** Selected gene ontology (GO) and Gene expression categories significantly enriched or underrepresented in the list of genes repressed in media with arginine at 30 min after antimycin A treatment. Analysis performed using online tool AnGeLi [60].

